# Mapping functional hemodynamic and metabolic responses to dementia: a broadband spectroscopy pilot study

**DOI:** 10.1101/2025.02.20.639231

**Authors:** Deepshikha Acharya, Emilia Butters, Alexander Caicedo, Li Su, John O’Brien, Gemma Bale

## Abstract

**Significance:** Broadband near-infrared spectroscopy (bNIRS) can simultaneously monitor several chromophores including the oxidative state of cytochrome c-oxidase (oxCCO), an oxygen metabolism biomarker whose activity is altered in Alzheimer’s disease. Being a portable and non-invasive neuromonitoring technique, bNIRS could thus provide accessibility to brain-specific biomarkers and aid in the dementia diagnostic pathway.

**Aim:** We use bNIRS recorded functional haemodynamic and oxCCO changes to assess their relevance in Alzheimer’s disease dementia as potential biomarkers.

**Approach:** Using a visual-stimulus paradigm, we recorded functional changes in oxy-, deoxy-haemoglobin and oxCCO in three similarly aged cohorts: healthy controls (n=5), individuals with mild cognitive impairment (n=7) and individuals with Alzheimer’s dementia (n=7). We then selected features from these functional responses to find the best correlation with clinical cognitive markers (cognitive and behavioural test scores and clinical diagnosis) using canonical correlation analysis.

**Results:** We found individual variations in peak amplitude and time-to-peak for all the stimulus-evoked bNIRS signals across the three cohorts. Canonical correlation analysis showed a strong correlation between bNIRS features and the clinical cognitive markers (r=0.902). However, repeating the same analysis by excluding the bNIRS oxCCO features leads to a significantly lower correlation (r=0.687) with the clinical markers.

**Conclusions:** oxCCO could be a crucial biomarker, partly explaining cognitive differences with dementia. bNIRS uniquely provides a portable and non-invasive technique to monitor several chromophores simultaneously including oxCCO with potential future applications in tracking dementia progression.

## 1 Introduction

Dementia arises from complex and heterogenous pathological conditions which lead to a general decline in cognition and impairments in daily function. There is huge variability in the clinical presentation of symptoms and underlying neuro-vascular degeneration not only in prodromal cases but also within specific dementia subtypes^1^. An early and differential diagnosis of dementia requires a holistic understanding of the presenting symptoms and their underlying causes. A combination of neuropsychological tests and neuroimaging techniques can help provide subject-specific comprehensive differential diagnosis. Neuropsychological tests assess several domains such as learning, retention, working memory, and motor function to quantify domain-specific cognitive decline^2^. In Alzheimer’s disease (AD), studies have consistently identified impairments in executive function and working memory. These cognitive deficits are also observed in the pre-clinical stage of mild cognitive impairment (MCI)^3^. Longitudinal monitoring using neuropsychological assessments, such as the Mini-Mental State Examination (MMSE)^4,5^ and tests evaluating executive dysfunction, can be instrumental in tracking the progression of AD. Concurrently, neuroimaging techniques such as magnetic resonance imaging (MRI) and positron emission tomography (PET) can provide structural, molecular and some functional insights into the underlying causes of the cognitive decline^6^.

In recent years, near-infrared spectroscopy (NIRS) has emerged as an alternative neuromonitoring technique in dementia research, bringing promise with its accessibility and portability relative to conventional neuroimaging methods^7^. NIRS uses the differential absorption of light in tissue in the near-infrared wavelengths (∼600-1200nm) to quantify changes in concentration of different tissue chromophores. Most common NIRS systems use sources operating at two wavelengths and measure the light received through a photodiode. This allows NIRS to resolve for two chromophores, most commonly, oxygenated (ΔHbO) and deoxygenated (ΔHbR) haemoglobin. During functional tasks, NIRS is often used to measure stimulus-evoked hyperaemia, characterised by an increase in local ΔHbO (and a decrease in ΔHbR) as oxygen is supplied to the cerebral region of activation. Previous NIRS studies have shown alteration in these functional hyperaemic signals in response to sensory stimuli with MCI and AD compared to healthy controls (HC), corroborating functional MRI findings^7^. In addition to oxygen supply, local oxygen metabolism is a critical indicator of brain health. Metabolic dysfunction is exaggerated in AD and may signal underlying issues such as neurovascular uncoupling and mitochondrial insufficiency^8^. Therefore, metabolic biomarkers have the potential to provide valuable insights into the pathophysiological pathways of AD.

Broadband NIRS (bNIRS) measures the differential absorption through tissue over a range of wavelengths simultaneously. Using a broadband light-source, such as a simple halogen bulb, paired with a spectrometer to detect the light received from the tissue, bNIRS can resolve for several chromophores concurrently. In addition to measuring changes in ΔHbO and ΔHbR, bNIRS can also measure concentration changes in the oxidation state of the mitochondrial enzyme cytochrome c-oxidase (CCO). Changes in the oxidation state of CCO (ΔoxCCO) represents changes in brain metabolism and has been validated in pre-clinical^9^, clinical^10,11^ and sensory stimulation^12–14^ research studies. While the relationship between ΔoxCCO concentration and AD pathology might not be linear, studies have shown that amyloid-β protein precursors reportedly block mitochondrial CCO and alternatively, this inhibition could instigate the shift towards the amyloidogenic pathway^15^. Post-mortem analyses have shown a 25-30% decrease in cortical CCO activity in AD patients which correlated with severity of cognitive impairment^16,17^. Meanwhile in at-risk groups with a maternal history of AD^18^ and subjects with MCI^19^, a decline in platelet mitochondrial CCO has been observed.

In this study, we use bNIRS to measure changes in local brain activity during visual sensory stimulation. We estimate functional changes in ΔHbO, ΔHbR and ΔoxCCO across healthy controls (HC), individuals with mild cognitive impairment (MCI), and those with Alzheimer’s disease (AD). Signal features such as peak amplitude and time-to-peak are used to calculate ΔHbO, ΔHbR and ΔoxCCO functional metrics. Haemoglobin difference (ΔHbD = ΔHbO - ΔHbR) is used as a proxy for cerebral blood flow^20^. Time lag between ΔHbD and ΔoxCCO during stimulus period is used as a proxy of neurovascular coupling, where an absence of the traditional ΔHbD lagging ΔoxCCO could indicate a mismatch in oxygen supply and demand. We analyse how these bNIRS signal metrics relate to the severity of cognitive impairment, with a particular emphasis on the role of ΔoxCCO-derived measures.

## 2 Materials and Methods

### 2.1 Study protocol

This study was conducted under the Optical Neuroimaging and Cognition protocol (IRAS project ID: 319284) sponsored by the University of Cambridge and the Cambridgeshire and Peterborough NHS Foundation Trust (CPFT). Participants expressing voluntary interest in the study who satisfied the eligibility criteria were provided with study related documentation and an official invitation. Home visits were scheduled, if convenient, for neuropsychological assessment and subsequently bNIRS recordings. Alternatively, participants underwent bNIRS recordings on site in the Department of Psychiatry at University of Cambridge. No difference in signal quality was found between at-home and on-site recordings. Individuals with severe dementia (MMSE<12), history of traumatic brain injury, history of excessive drug or alcohol use, conditions affecting haemodynamics and metabolism or significant physical or psychiatric illnesses were excluded from the study.

Upon consent, each participant was scheduled for an at-home neuropsychological assessment, testing across a range of cognitive functions. For each participant, a cumulative executive function test (ExFT) score was calculated from their individual scores in test for inference sensitivity (conflicting instructions), inhibitory control (go-no go) and digit span. Additionally, participants underwent the Mini Mental State Examination (MMSE)^4^, and these scores were incorporated into the subsequent analysis. Any participant that did not complete MMSE or ExFT, was not included in further analysis. ExFT was scored out of 19 and MMSE was scored out of 30. In both cases, lower scores indicated more cognitive impairment. Statistical testing for inter-group differences in the scores was done using a Wilcoxon rank sum test (*ranksum*, MATLAB 2023a). bNIRS recordings were a small pilot portion of the larger study trying to understand the role of NIRS, specifically high-density NIRS, in dementia diagnostics.

Measurements were performed using a miniature bNIRS system^21^ with the Ocean Optics HL2000 white light source and a long-pass filter with a cut-on wavelength of 630nm (Thorlabs, FGL630) to limit input light to near-infrared range. A custom Wasatch spectrometer (WP-VISNIRX-C-S-25) configured for 650-910nm was used to detect the light received from the tissue. Custom 90° flat optical fibres (Engionic Fiber Optics GmbH) were used to interface devices (source and detector) to the participant. An optode holder with a source-detector separation of 3cm was 3D printed from flexible resin to have a comfortable probe design and provide maximal contact to participant head. The probe was attached to participant head using a fabric band.

The probe was placed over the right visual cortex while a full-field checkerboard stimulus was presented to the participant. A radial checkerboard stimulus, reversing at 7.5Hz for 10 seconds was repeated 12 times with an inter-block rest period of 15 seconds, jittered by 0.1 second. The stimulus was implemented in Python using PsychoPy (v 2021.2.3)^22^. A schematic representation of the experiment structure is shown in Figure 1a.

**Fig. 1.**
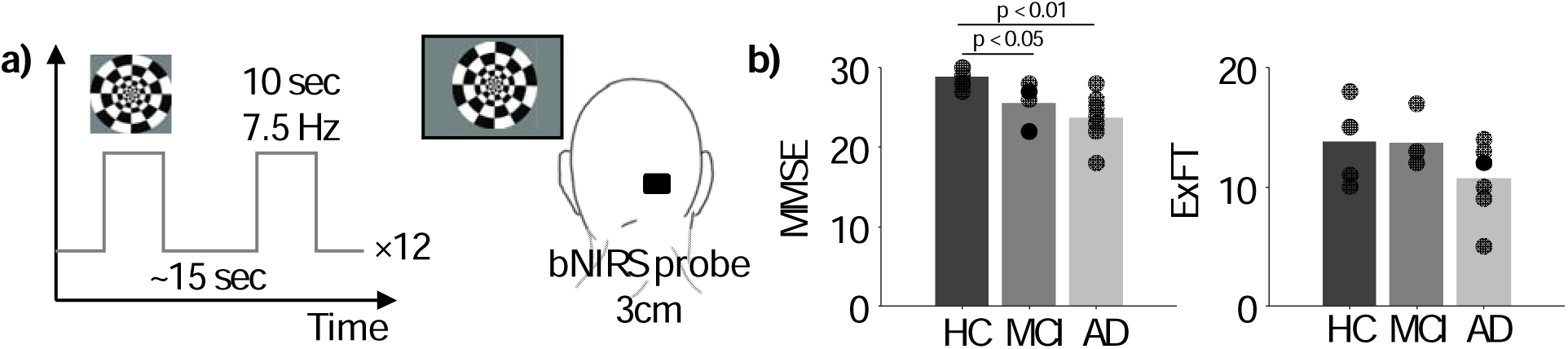
Experimental setup and cohort: (a) visual stimulus protocol and the bNIRS setup and (b) neuropsychological test scores for the 3 cohorts based on clinical diagnosis. Each bar shows the mean score and the points overlaid show individual scores for each cohort. MMSE was scored out of a maximum of 30 and ExFT out of 19.

### 2.2 Data Preprocessing

Participants’ bNIRS data was rejected if the maximum spectral intensity of the received light was less than 1000 or if the participant lacked a distinct stimulus-locked peak in the Fourier transform of the bNIRS data. Additionally, through visual inspection, if a participant’s absorption spectra were found to be noisy, the participant was rejected from further analysis. The remaining participants’ bNIRS data was then pre-processed for further analysis. All signal analysis was done using MATLB R2023a.

Noise in spectral data was identified by spikes on the absorption spectra crossing a z-score threshold of ±5, potentially arising from contamination with ambient light. Timepoints corresponding to the noisy spectra were noted. The calculated changes in concentration values at those timepoints removed and the missing data filled in using spline interpolation. The spectral data was converted to changes in concentration (ΔHbO, ΔHbR and ΔoxCCO) using the UCLn algorithm, a generalised modified Beer-Lambert Law algorithm^23^. The data was re-sampled to 2Hz and a wavelet motion correction^24^ with an inter-quartile range of 1.5 (adapted from Homer3 toolbox^25^) was applied to remove motion spikes from the changes in concentration timeseries. The resulting signals were filtered using a 3^rd^ order Butterworth low pass filter (0.08Hz cutoff frequency) in series with a 5^th^ order Butterworth high pass filter (0.01Hz cutoff frequency) to remove low frequency noise and physiological signals such as Mayer waves, respiration, and heart rate.

### 2.3 Functional bNIRS Responses and Feature Selection

Stimulus-evoked functional responses and associated features were calculated for ΔHbO, ΔHbR and ΔoxCCO respectively. At the beginning of each stimulus block, a manual marker was input by the experimenter which was used to epoch each timeseries. For each signal epoch, the average of 3 seconds before stimulus onset was used for baseline correction. Each baseline corrected signal epoch included 10 seconds of stimulus presentation followed by 15 seconds of baseline, for a total block duration of 25 seconds. Epochs with responses whose peak was within ± 3 standard deviations from the mean of the pre-stimulus period were rejected due to lack of a significant response. The remaining epochs were averaged to give a single evoked ΔHbO, ΔHbR and ΔoxCCO epoch per participant.

Two features were selected to quantify these evoked responses for further analysis. The widest prominent peak from the average evoked signal was identified (*findpeaks*, MATLAB) as the haemodynamic response. The corresponding peak was adjudged the peak amplitude (PA) and the corresponding time was the time-to-peak (TTP).

Additionally, cross-correlation between haemoglobin difference (ΔHbD) and ΔoxCCO timeseries was calculated with a max-lag threshold of ±12.5 seconds (spanning 25 seconds of the block duration). Lag at maximum absolute correlation was used to quantify time-lag between the two signals.

### 2.4 Canonical Correlation Analysis

Canonical Correlation Analysis (CCA) is a statistical method used to assess multivariate relationships. This technique is especially useful when working with intercorrelated variables which is often the case in complex disease classification and neuroscience^26^. To apply CCA to our analysis, two variable sets were created – bNIRS metrics and cognitive metrics. The bNIRS metrics comprised PA and TTP for ΔHbO, ΔHbR and ΔoxCCO each, and time-lag between ΔHbD and ΔCCO. Meanwhile, the cognitive metrics comprised of the MMSE and ExFT scores, and a cohort code based on their clinical diagnosis (1 for AD, 2 for MCI and 3 for HC). CCA was used to identify the relationship between bNIRS and cognitive metrics and assess the amount of variance in one, explained by the other set. Additionally, associated loadings to each of the variables helped gauge their role in this prediction. Non-parametric statistical testing was done by creating a null set^26^ where cognitive metrics were randomly drawn from their set of possible values (1-30 for MMSE, 1-19 for ExFT and 1-3 for the cohort code) and the resulting set used to calculate CCA with the original bNIRS metrics. This was repeated 5000 times to create a distribution of canonical correlations with a null cognitive set. Significance was tested by comparing the original canonical correlation value to the 90^th^ percentile of this null distribution.

To specifically test the importance of ΔoxCCO in this analysis, CCA was repeated excluding the ΔoxCCO-related signal metrics (PA and TTP of ΔoxCCO and time-lag). The resulting canonical correlation was then compared to the initial findings using bootstrap analysis. Bootstrapping was done by randomly drawing paired samples from the canonical variate set 5000 times, with replacement, and calculating the corresponding correlations to create a distribution of correlation values. This was done for each test case (with and without ΔoxCCO) and the resulting distribution of correlations were compared to test for significant differences using 2-tailed t-test.

## 3 Results

Data was collected from 56 participants (HC=23, MCI=19 and AD=18). After applying the data-rejection criteria outlined earlier, we rejected 8 participants each due to low intensity of received light and noisy recorded spectra. Additionally, 21 participants were rejected due to the lack of a reliable stimulus-locked response in the bNIRS recorded concentration signals. The extremely high attrition rate may be attributed to factors such as poor contact of the optical probes due to thickness or colour of hair, noise due to motion or ambient light, or lack of participant engagement during the passive-stimulus task. We address potential solutions in the Discussion.

Here we report data across 3 similarly aged groups of – HC (n=5, age: 72.72±9.57 years), MCI (n=7, age: 78.14±5.3 years) and AD (n=7, age: 78.4±7.57 years).

The average and individual scores for MMSE and ExFT across the 3 cohorts are plotted in Figure 1b. As expected, scores for both cognitive tests were the lowest for the AD group (MMSE: 23.7, ExFT: 10.7), followed by MCI (MMSE: 26.1, ExFT: 12.6) and the highest for HC (MMSE: 28.8, ExFT: 13.8). MMSE scores for MCI and AD were significantly lower than HC (p-value_MCI-HC_ = 0.02, p-value_AD-HC_ = 0.007). ExFT scores showed no significant group differences.

### 3.1 Functional bNIRS Responses and Feature Differences Between Cohorts

As expected, in response to visual stimuli, we observed increases in ΔHbO and ΔoxCCO and a decrease in ΔHbR across all three cohorts. Figure 2a shows the average functional epochs for the different bNIRS signals for HC, MCI and AD groups. Average participant responses were grouped based on their clinical diagnosis. Average PA for ΔHbO haemodynamic responses was the highest in the AD group (0.18±0.07µM) followed by HC (0.11±0.06µM) and the lowest for MCI (0.06±0.06µM). A similar trend was observed for ΔoxCCO (HC = 0.04±0.01µM, MCI = 0.03±0.01µM and AD = 0.05±0.02µM, respectively) and ΔHbR (HC = -0.07±0.03µM, MCI = - 0.08±0.03µM and AD = -0.03±0.02µM, respectively). Further, on average, an early TTP was seen in the MCI group for ΔHbO (11.4±1.45s) and ΔoxCCO (10.7±1.55s) responses compared to AD (ΔHbO = 12.1±1.39s ΔoxCCO = 12.2±1.73s) and HC (ΔHbO = 13.2±1.59s ΔoxCCO = 13.6±1.23s). The opposite trend was observed for ΔHbR with TTP on-average being earlier for HC (11.7±1.28s) compared to MCI (13.6±0.58s) and AD (12.2±1.19s). Finally, ΔHbD was leading ΔoxCCO for MCI (1.7±1.44s) and AD (1.07±1.87s) but lagging ΔoxCCO for HC (2.3±2.6s). Figure 2b-d shows the average ± standard error for PA, TTP and lag respectively for each group. It must be noted here that all values are reported as mean ± standard-error and none of the metrics showed significant difference between groups.

**Fig. 2.**
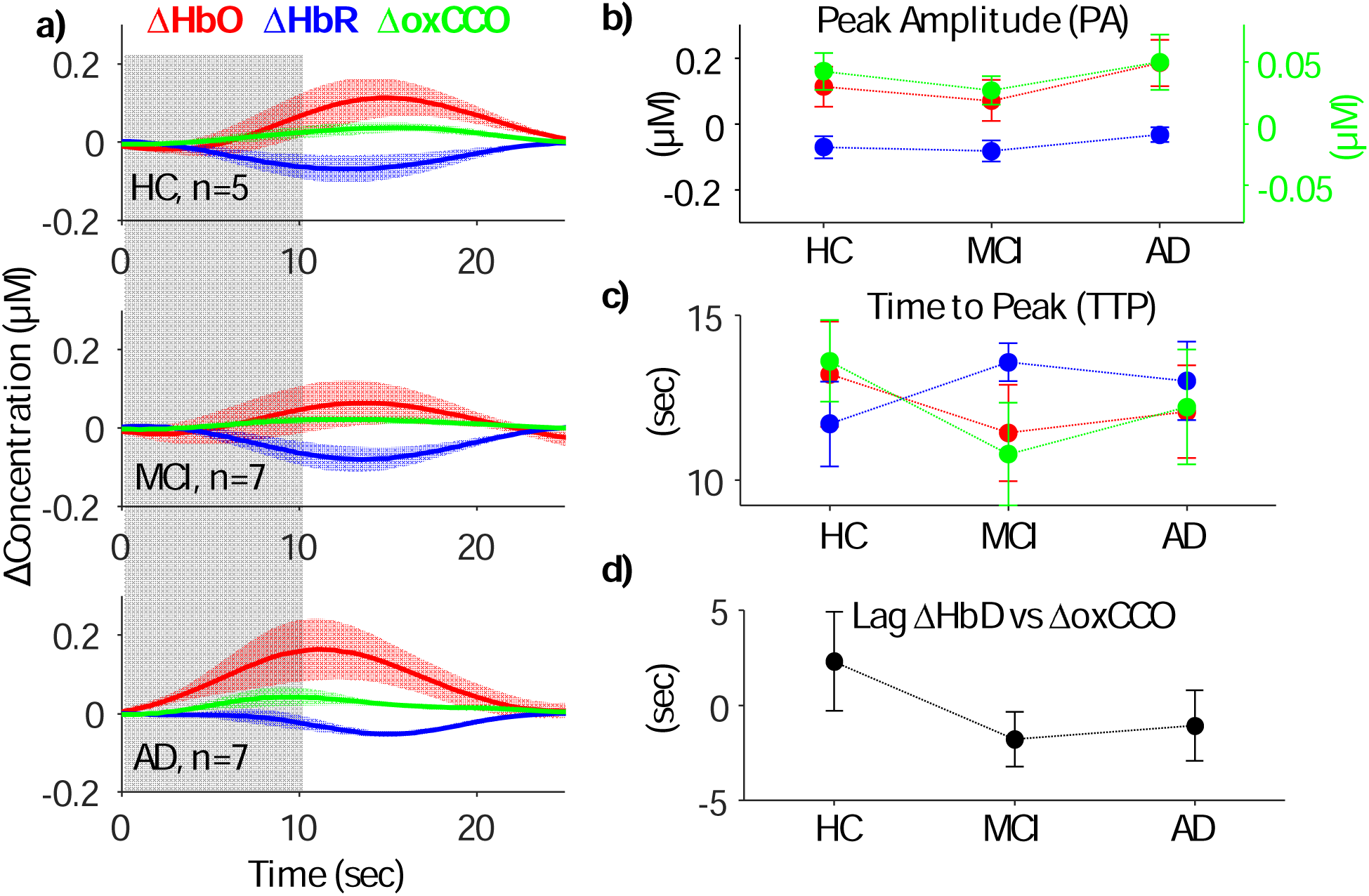
Functional responses to visual stimulation recorded by bNIRS: a) changes in concentration of ΔHbO, ΔHbR and ΔoxCCO epoched at the stimulus onset to 25s (10s stimulus block (gray) + 15s rest). For each group, the bold line shows the average response within the group and the shaded region shows the standard error around the mean. b-d) shows the PA, TTP and lag for each group. The filled circles show the mean and the lines around the standard error. For the entire figure, values associated with ΔHbO are in red, ΔHbR in blue and ΔoxCCO in green.

### 3.2 bNIRS and Cognitive Metric Correlations

A Pearson’s correlation was calculated between the bNIRS metrics (PA and TTP for ΔHbO, ΔHbR and ΔoxCCO and time-lag) and the cognitive metrics (MMSE and ExFT scores and cohort code) to assess how they each contributed to explaining the variance in the other metric. Data from all participants was pooled for this analysis. ExFT scores showed the strongest correlation with all the bNIRS metrics, specifically PA and TTP for ΔHbO and ΔoxCCO. Figure 3a shows the correlation between each set of variables. The same pooled dataset across all participants from above was used to perform CCA between the bNIRS and the cognitive metrics. The highest correlation corresponding to the first component of the bNIRS and cognitive metrics was 0.902. Loadings were also calculated from the first components to understand their relationship to the corresponding metrics. Figure 3b shows the loadings for each metric and the highest canonical correlation between the two variable sets.

**Fig. 3.**
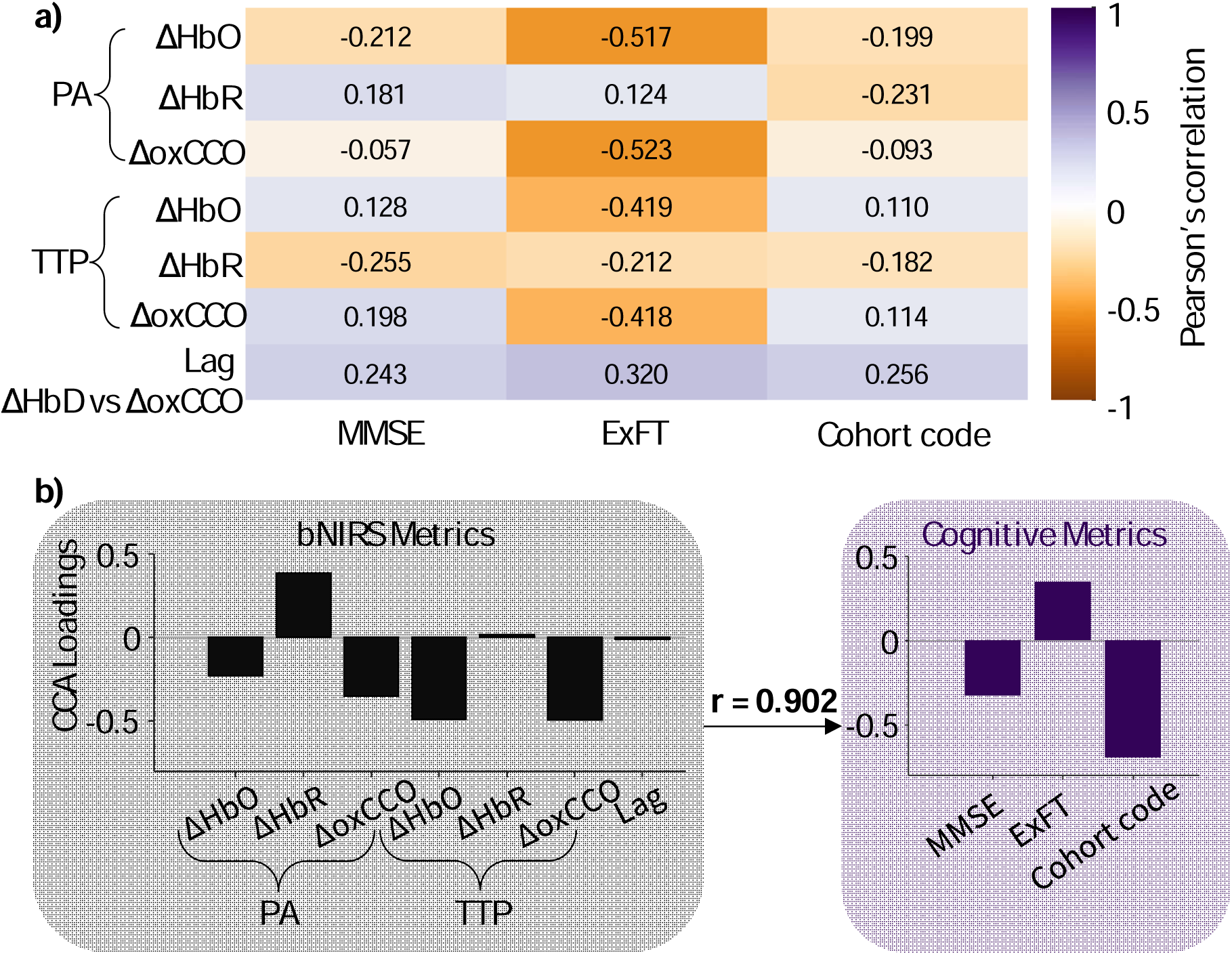
Correlations between bNIRS and cognitive metrics: a) Pearson’s correlation between the two sets of metrics. b) CCA results between the two sets of metrics – the bar graphs represent the corresponding loadings for each variable. All the bNIRS metrics are shown in black and the cognitive metrics are shown in purple. The two sets were found to have canonical correlation of 0.902. This value was found to be greater than the 90^th^ percentile of the null distribution of canonical correlations.

### 3.3 Contribution of ΔoxCCO in Clinical Metric Correlations

Canonical loadings for ΔoxCCO PA (-0.35) and TTP (-0.49) were found to be amongst the highest for the bNIRS metrics, though time-lag was a low (-0.01). We further tested the importance of ΔoxCCO in accounting for the variance in the clinical data by performing CCA once including ΔoxCCO and once without. When excluding ΔoxCCO, the new bNIRS metric set only used PA and TTP for ΔHbO and ΔHbR. Figure 4a shows the highest correlations between the first components for each of the test case - with ΔoxCCO (r = 0.902) and without ΔoxCCO (r = 0.687). To test the significance of the difference in the correlations, we also performed a bootstrap analysis followed by a 2-tailed t-test. Figure 4b shows the distribution and the 10^th^ and 90^th^ percentile of each distribution. The CCA correlation when using ΔoxCCO was found to be significantly higher compared to without ΔoxCCO (p-value<<0.05).

**Fig. 4.**
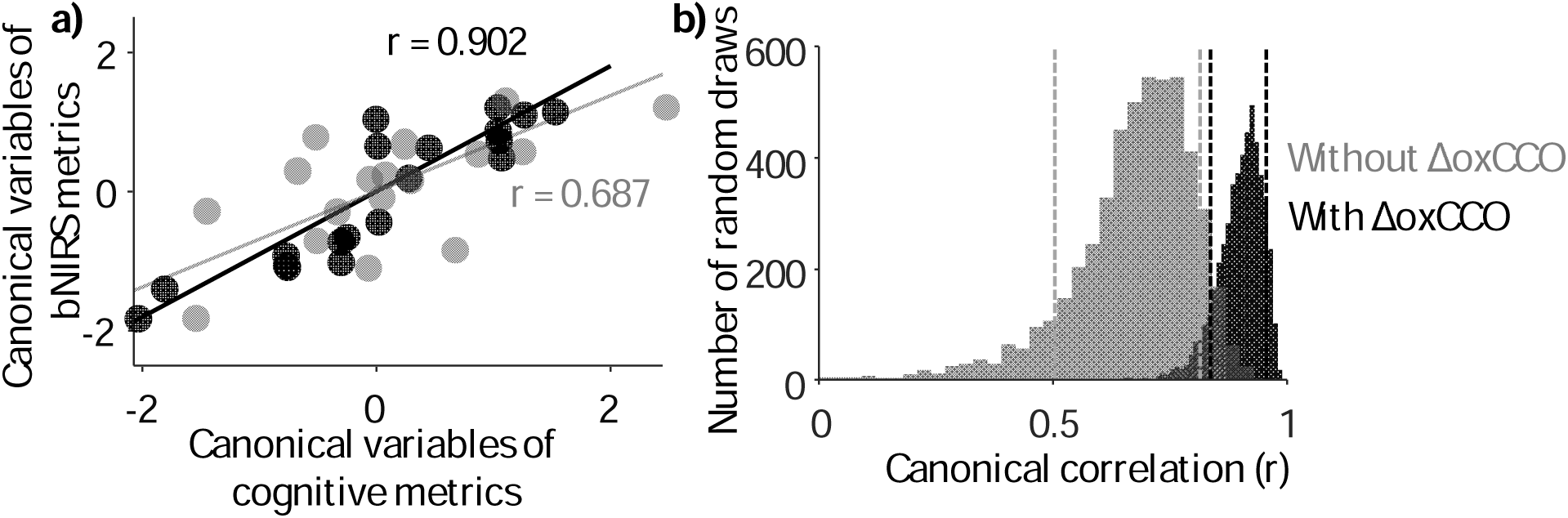
CCA for two cases – bNIRS metrics with ΔCCO (black) and bNIRS metrics without ΔCCO (gray): a) the CCA correlation between the canonical variates for the two cases, b) The bootstrap distribution of correlation values for the two cases. The dashes vertical lines show the 10th and 90th percentile of the respective distributions.

## 4 Discussion

In this work, we propose the use of bNIRS for the first time in dementia research, leveraging its ability to resolve multiple biomarkers simultaneously. We specifically focus on the role of ΔoxCCO, a key mitochondrial enzyme, in AD. Using bNIRS in a visual-stimulus paradigm, we estimate visual-evoked haemodynamic (ΔHbO and ΔHbR) and metabolic (ΔoxCCO) changes across 3 similarly-aged cohorts – AD (n=7), MCI (n=7) and HC (n=5). To comprehensively understand the relationship between the recorded signals and diagnosis, we distilled the results in 2 variable sets, bNIRS metrics (PA, TTP and time-lag) and cognitive metrics (MMSE and ExFT scores and the cohort code) and used CCA to find the correlation (r = 0.902). Applying the same canonical analysis without the ΔoxCCO-derived metrics yielded in a significantly lower (p-value<<0.05) canonical correlation (r = 0.687). This section discusses our results in the context of existing literature and suggests improvements for translation of this work.

We observed an expected decrease in average neuropsychological test scores (Figure 1b) from HC to AD, for both MMSE and ExFT. MMSE test scores were significantly lower (p-value<<0.05) for MCI and AD compared to HC. Meanwhile ExFT scores showed no significant difference between groups. The variance in individual scores was higher within groups for ExFT compared to MMSE. Previous work has linked this heterogeneity in executive dysfunction in mild AD to potential differences in underlying disease processes and secondary conditions^27^. Figure 1b shows individual scores overlaid on the corresponding bar plot showing group averages. During functional bNIRS recordings, we also observed expected functional hyperaemic responses though their PA and TTP varied across the three groups (Figure 2). Neurovascular dysfunction has been widely reported in AD corroborated by the hypoactivation reported during sensory stimulation/tasks with increasing cognitive decline^28^. Consequently, we did observe a decrease in PA and a faster TTP in MCI compared to HC for ΔHbO. This reduction in occipital activation has been previously reported during a working-memory task^29^ in MCI compared to HC groups. Further, Bu et al^30^ also showed a reduction in coupling strength at resting-state across different brain regions, including the occipital lobe, in MCI compared to HC. While the MCI group showed reduced activation amplitude compared to HC, we observed the opposite for the AD group. An average increase in PA was seen between HC and AD for ΔHbO. This hyperactivation in the visual cortex in early-stage AD has been seen in previous studies and linked to compensatory mechanisms or dysregulation of the neurovascular homeostatic cycle^31,32^. This could also explain the faster TTP in MCI and AD compared to HC for ΔHbO (Figure 2c). Interestingly, the difference in TTP between ΔHbO and ΔHbR was the highest for MCI which could originate from systemic sources or cortical haemodynamic differences^33^ amplified by the neuro-vascular deficiencies with dementia. This difference in TTP was however absent for AD.

Further investigation by potentially isolating systemic effects with short channel regression might shed light on differences in TTP between the haemodynamic signals with dementia. We also observed a decrease in ΔHbR PA response from HC to AD despite an increase in ΔHbO PA. Metabolic differences with AD could partially explain this discrepancy.

CCO is an important part of the oxidative metabolism of glucose featuring in the electron transport chain. Changes in absolute concentrations of complex IV (CCO) have been related to AD. In a post-mortem study relating CCO changes with AD, they found the expected decrease in CCO in the frontal and parietal regions, however a non-significant elevation in CCO in the occipital cortex^34^. Contrarily, another study^16^ found a 25-30% decrease in CCO across all cortical regions studied, including the occipital cortex. Beyond absolute CCO concentration, in AD mouse models, a deficient oxygen consumption at complex IV in the mitochondria has been observed^35,36^ and could specifically explain reduced concentrations of oxCCO. While most measurements of CCO are postmortem in the cases of dementia, measurements of glucose metabolism with FDG-PET have enabled similar assessment of regional cortical metabolic deficiencies during disease progression. Condition specific hypometabolism have been reported in AD^37^ and mapped to eventual disease manifestation in MCI^38^. During functional tasks, a study found reductions in cerebral metabolic rate of glucose in AD compared to HC during a visual recognition task using FDG-PET^39^. We also observed a small decrease in functional ΔoxCCO PA in the visual cortex for MCI group compared to HC (Figure 2). However, ΔoxCCO PA was highest was AD, potentially indicating hypermetabolism. Another common measure of metabolism in MRI studies is cerebral metabolic rate of oxygen consumption (CMRO_2_). A study showed reduction in frontal cortical blood flow and CMRO_2_ as well as their correlation (coupling) in participants with subjective cognitive decline compared to controls^40^. Beside differences in correlation flow-metabolism coupling is also seen with differences in time lag between the two signals. We found flow (ΔHbD) to lag metabolic changes (ΔoxCCO) in HC by a few seconds which is expected during functional activation in a healthy brain due to neurovascular coupling^41^. However, this was reversed in MCI and AD where ΔHbD was found to lead ΔoxCCO. No differences were observed in the correlation between the signals at these lag times. The delayed metabolic response compared to flow and the faster TTP of ΔoxCCO and ΔHbO compared to HC could indicate underlying neuronal deficits, impaired cerebrovascular reactivity, or potential neurovascular uncoupling. It must also be noted here that ΔoxCCO may not match CMRO_2_ under some conditions^42–44^, which might affect the interpretation of our results against the few existing functional activation dementia literature that usually use CMRO_2_. AD has been dubbed as a metabolic disease^45^ with markers such as oxCCO potentially being key to aiding diagnosis. Our work here is the first to bring the more established mitochondrial marker (oxCCO) for AD to non-invasive functional studies to be able to assess differences in PA, TTP and correlation with other haemodynamic signals with dementia. Further investigation of these results with different sensory tasks, a true measure of local blood flow and a simultaneous assessment of CMRO_2_ would help better understand the results in the context of the existing literature. Finally, while there were trends between groups for the bNIRS metrics, none of the trends were found significant. One possible explanation for this could be the within-group heterogeneity in prodromal and early-AD groups arising from heterogeneity in underlying disease pathways and differences in disease progression.

All the collected data was pooled into two variable sets – bNIRS metrics and cognitive metrics and a pairwise Pearson’s correlation calculated. The strongest correlation was found between the ΔHbO and ΔoxCCO PA and TTP and the ExFT (Figure 3a). Recently more studies have indicated early impairment of executive function in AD, possibly arising from degeneration of prefrontal cortex^46^. However, given the interdependence of variables in each set, conclusions from pairwise Pearson’s correlation might be conflated. For a more comprehensive understanding, we used CCA, a multivariate statistical analysis method. With CCA we found that the first component of bNIRS metrics accounted for 80% of the variance in the first component of the cognitive metrics (Figure 3b) with a correlation of r=0.902. This correlation was found to be greater than the 90^th^ percentile of the null canonical correlation distribution created by shuffling a randomly drawn set of cognitive metrics. However, the canonical correlation significantly decreased (p-value <<0.05) to r=0.687 when not using ΔCCO-derived metrics, now accounting for only 50% of the variance in cognitive metrics (Figure 4). This highlights the crucial role bNIRS recorded ΔoxCCO could play in understanding differences in cognitive impairment with AD. This was supported by the correspondingly high loadings for the ΔoxCCO features, particularly PA and TTP, which represent its high contribution to the final correlation value (Figure 3b). We also observed a similarly high contribution from ΔHbO and an expected inverse contribution from ΔHbR given its negative magnitude for PA and opposite trend in TTP compared to ΔHbO and ΔoxCCO (Figure 3b). Overall large loadings of the cognitive metrics indicate the role of all of them – MMSE, ExFT and cohort code based on diagnosis, in the relationship with bNIRS metrics. Finally, our CCA results indicate the importance of using multiple biomarkers, especially metabolic markers such as ΔoxCCO to improve differential diagnosis.

Crucially, it must be noted that the functional bNIRS data was collected at-home or at a controlled-environment study site, based on participant convenience. No data quality differences were found between the two study locations. Our work is amongst the first to use bNIRS for AD, especially at-home. This highlights the importance of portable and non-invasive neuromonitoring techniques such as bNIRS in enabling at-home monitoring of symptoms and aiding dementia diagnosis.

The results here are from a small cohort of participants using a single channel bNIRS setup during a visual task. Due to challenges in data collection, despite the 56 complete datasets of recorded participants, only 19 were used for final analysis. This led to 3 cohorts with small sample sizes that were not perfectly age- or sex-matched. Influence of age and sex on vasculature could impact the final results. Future improvements to this work would include a greater number of bNIRS channels to have a better coverage of the sensory region, improved signal-to-noise ratio, and removal of the influences from superficial layers such as scalp. Additionally, a high-density multi-channel setup could enable cortical source localisation^47^ which concurrently with patient-specific MRI could help account for signal difference due to brain atrophy, especially in advanced AD cases. Another aspect of study improvement could be in the task itself. Given the differences in cortical vulnerabilities with AD^48^, a battery of functional tests probing different cortical regions could improve dementia/severity classification as well as participant engagement. Correspondingly, the bNIRS probe setup could be moved to different regions to acquire region specific haemodynamic responses to the task. While conventional CCA can be very informative for small datasets, it is also heavily affected by outliers which made signal quality checks even more imperative for our analysis. Effect of outliers can be limited with better signal quality, for which we have discussed recommendations above, or with larger datasets. With larger datasets, it will be feasible to develop a classifier using machine learning algorithms and provide a more robust understanding of the role of ΔoxCCO in aiding AD diagnosis.

## 6. Conclusion

Neuropsychological and neuromonitoring techniques could provide early and differential diagnosis of dementia and enable targeted symptom management. Metabolic dysfunction, specifically CCO, could play a key role in understanding early cognitive impairment arising from AD. bNIRS provides a portable and non-invasive alternative to monitoring haemodynamic and metabolic changes and could aid dementia diagnosis. Here, in a pilot study, we used bNIRS to record concentration changes in ΔHbO, ΔHbR and ΔoxCCO during visual stimulus presentation in participants with MCI, AD and HCs. Using CCA, we showed a strong correlation between the bNIRS signal features and the cognitive metrics, which significantly reduced when excluding ΔoxCCO-derived metrics. Thus, bNIRS recorded metabolism (ΔoxCCO) has potential in aiding dementia diagnostics and could also aid regular monitoring of treatment outcome due its compatibility to use at-home.

### Disclosures

John O’Brien has acted as a consultant for TauRx, Novo Nordisk, Biogen, Roche, Lilly, GE Healthcare and Okwin and received grants or academic in-kind support from Avid/Lilly, Merck, UCB and Alliance Medical. He is supported by the NIHR Cambridge Biomedical Research Centre and the Medical Research Council funded Dementias Platform UK.

### Code, Data, and Materials

Relevant code will be shared on final publication on Code Ocean.

## Acknowledgments

We would like to thank the participants, their carers and families for volunteering for the study and specially acknowledge Emilia Butters for undertaking all the data collection. We would also like to acknowledge the help from Dr. Elizabeth McKiernan, Dr. Peter Swann and Dr. Carlos Muñoz-Neira for help with participant recruitment. Gemma Bale and Deepshikha Acharya would like to thank the Newton Trust for sponsoring a part of this research. Gemma Bale and Emilia Butters acknowledge funding from the Gianna Angelopoulos Programme for Science and Technology Innovation.

